# Broad and Long-lasting Immune Response against SARS-CoV-2 Omicron and Other Variants by PIKA-Adjuvanted Recombinant SARS-CoV-2 Spike (S) Protein Subunit Vaccine (YS-SC2-010)

**DOI:** 10.1101/2021.12.22.473615

**Authors:** Yuan Liu, Nan Zhang, Bin Wang, Yi Zhang

## Abstract

Recently SARS-CoV-2 Omicron (B.1.1.529) variant was identified in South Africa with numerous mutations in spike protein, and numerous community infections have been reported and raised grave concern around the world. Some studies found that the neutralization effects of several licensed vaccines against Omicron were dramatically reduced, which significantly affected antibody mediated protection, especially for individuals whose immunization were completed after extended period. In this regard, we studied the persistence and neutralization activity toward mutant strains in animal serum immunized with PIKA-adjuvanted recombinant SARS-CoV-2 spike protein subunit vaccine (YS-SC2-010). Here we are reporting that animal serum collected at 596 days after immunization with YS-SC2-010 still retains high and persistent neutralizing activity against all the Variant of Concern (VOC) variants, including Omicron variant. Although it is a blessed event to achieve 20 months long neutralization against Omicron variant after immunization with YS-SC2-010, it was also founded that the neutralization effect of immune serum on Omicron decreased by 6.29 folds as compared to D614G, more significantly when compared with other mutant strains.

---

On November 9, 2021, SARS CoV-2 variant Omicron was first reported in South Africa composed of numerous mutations at the spike protein unit, which has become the main antigen target for vaccine design^1^. Therefore, the emergence of Omicron has aroused a new round of concerns about the effectiveness of existing vaccine and the risk of reinfection. Compared with the wild-type strain, S protein of Omicron has 30 non-synonymous substitutions, three small deletions and one insertion. Among them, 15 mutations are located in the receptor binding domain (RBD), which is the main target of neutralizing antibodies^2^. Several S protein mutations observed in Omicron have also appeared in previously mutated strains, such as alpha, beta, gamma, delta, kappa, zeta, lambda and mu, which may lead to higher transmissibility and immune escape^3,4^. Since there are multiple mutation sites in Omicron S protein, which may produce aggregate impact on reducing the efficacy of current vaccines, it is urgent to investigate the neutralization effect of the vaccine on Omicron to judge whether it can prevent Omicron infection. Some studies found that the neutralization effects of vaccines on Omicron were seriously reduced, which significantly affected antibody mediated neutralization, especially for individuals immunized for a long time^5–7^.

We have previously published the long-term humoral immune response induced by PIKA-adjuvanted recombinant SARS-CoV-2 spike (S) protein subunit vaccine (YS-SC2-010) from 14 days to 406 days, and the results showed that the neutralizing antibody still retain with impressive neutralization property one year after immunization^8^. 406 days after the first immunization, the pseudovirus neutralizing antibody level gradually decreased from 10^4^ to 10^3^, indicating that the vaccine can achieve long-term humoral protection over 13 months.

To further study the durability of immune response, our observation period spanned more than 590 days. The neutralizing antibodies in sera collected at days 448, 518 and 596 after the first immunization were detected (Fig 1F). Our studies showed that all serum samples were able to neutralize SARS-CoV-2 pseudotyped viruses including the spike protein of the wild-type strain, and several circulating variants, including D614G, B.1.1.7 (Alpha), B.1.351 (Beta), P.1 (Gamma), B.1.617.2 (Delta) and B.1.1.529 (Omicron). The mean neutralization titer (50% effective dilution, ED50) of D614G, Alpha, Beta, Gamma, Delta, and Omicron at day 596 were 1335.72, 765.47, 380.94, 620.56, 1207.39, 140.11 respectively. The results demonstrated that the neutralizing antibody against variant strains could still be positive and maintained at a stable level, indicating that the vaccine could provide long-term humoral protection for 20 months.

**Fig.1.**
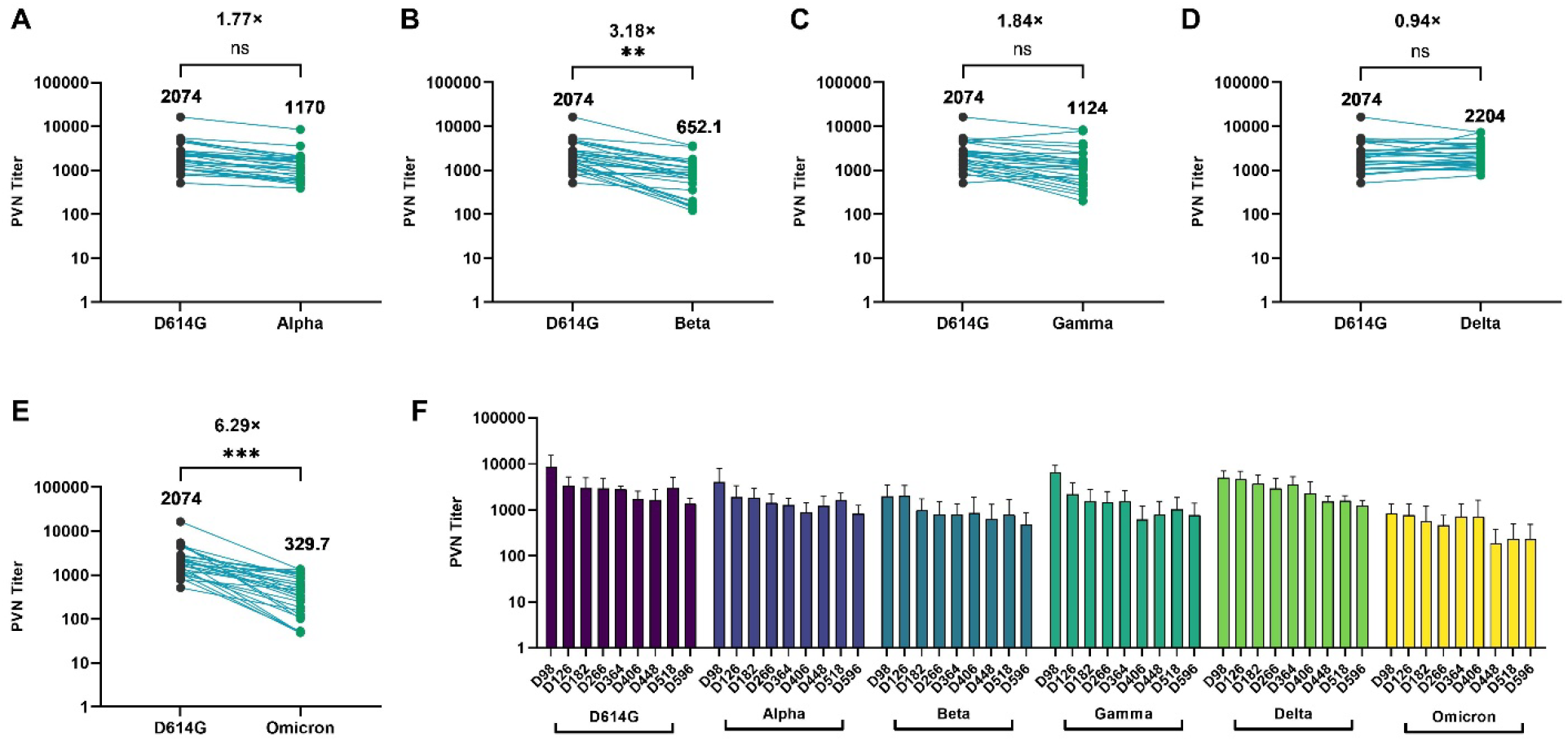
Antibody-mediated neutralization efficacy against pseudotyped SARS-CoV-2 variants. Rabbits were injected intramuscularly with YS-SC2-010 at 0, 7 and 14 days, and pseudovirus neutralization antibody levels at 98, 126, 182, 280, 364, 406, 448, 518 and 596 days after the first immunization were determined using the recombinant VSV-based SARS-CoV-2 pseudovirus. (A-E) The neutralization ED50 and ratio compared to the reference strain D614G was displayed as indicated. For all panels, values above the symbols denote geometric mean titer (n=28, *P*-values were analyzed with unpaired t-test, \*\**p* < 0.01, \*\*\**p* < 0.005, ns not significant). (F) Kinetics of pseudotyped SARS-CoV-2 variants neutralizing antibodies over 596 days after vaccination.

To test the potential neutralizing response against different variants induced by YS-SC2-010, sera from the rabbits vaccinated with YS-SC2-010 were obtained on certain days after prime immunization. We assayed the neutralization activity against above five SARS-CoV-2 pseudotyped viruses. The mean neutralization titer (ED50) of Alpha, Beta and Gamma is reduced about 1.77, 3.18, and 1.84-fold respectively from the baseline reference strain D614G (Fig. A-C), while the neutralization for Delta variants is higher than the reference strain D614G (Fig. D). As for Omicron, which is the most concerned new strain all over the world, the ED50 was 329.7, which represented about 6.29-fold reduction of neutralization compared to the reference strain D614G (ED50=2074) (Fig. 1E).

Various preclinical and clinical studies showed that neutralizing and binding antibody titer is correlated to the protection efficacy of vaccine^9–11^. The published research data on Omicron found that the Omicron variant had the most significant escape from the serum of subjects vaccinated with COVID-19 vaccine^6,7,12^. Compared with Delta strain, the antibody titer against Omicron decreased by more than 90%^12^. Our study also found that the neutralizing antibody in the serum of animals inoculated with YS-SC2-010 was indeed lower than other variants in neutralizing Omicron variant. However, despite the escape of Omicron from the immune serum, the neutralizing antibody against Omicron in the immune serum 596 days after the vaccination remained at a high level. It is suggested that the vaccine is still expected to be effective in preventing Omicron. At present, YS-SC2-010 is in clinical trial stage, and more human data will be published in the near future.

## Materials and methods

### Vaccine formulation

The YS-SC2-010 formulations were prepared with concentrations of 6 μg/ml for S-trimer protein and 1 mg/ml for the PIKA adjuvant. The vaccine is produced according to Good Manufacturing Practices by Liaoning Yisheng Biopharma Co., Ltd.

### Immunogenicity analysis of S-Trimer in rabbits

New Zealand rabbits were immunized intramuscularly with YS-SC2-010 three times on Day 0, Day 7, and Day 14. Serum were collected from each rabbit in each group on day 98, 112, 126, 154, 182, 238, 322, 364, 406, 448, 518 and 596 after the initial immunization and tested for pseudotyped SARS-CoV-2 neutralizing antibodies to evaluate the immune persistence.

### Pseudo-virus neutralization assay

SARS-CoV-2 (D614G, Alpha, Beta, Gamma, Delta, and Omicron) pseudo-virus neutralization assays were performed at Gobond Testing Technology (Beijing) Co., Ltd. (Beijing, China). Vero cells were cultured in DMEM supplemented with 10% heat inactivated fetal bovine serum, 50 U/ml Penicillin-streptomycin solution at 37°C with 5% CO_2_. Inactivated serum samples were serially dilute and incubated with 1.3×10^4^ TCID50/ml SARS-CoV-2 pseudo-typed virus for 1 h at 37°C. Vero cells were added after 1h and allowed to incubate for 24 h. Positive and negative control samples were prepared as same way. Post infection, cells were lysed and RLU were measured using the Microplate Luminometer. Neutralization titers were calculated as the serum dilution at which RLU were reduced by 50% compared with RLU in virus control wells.

### Statistical analysis

Statistical analysis was performed using Prism 9.0 (GraphPad software). The two groups were compared using an unpaired Student *t*-test. P values < 0.05 is considered to be significant. * *p* < 0.05, \*\**p* < 0.01, *** *p* < 0.001, \*\*\*\**p* < 0.0001. NS, not significant.

## Acknowledgements

The authors would like to acknowledge and thank Gobond Testing Technology (Beijing) Co., Ltd. for providing the SARS-CoV-2 pseudotyped viruses and conducting the neutralization test.

## Ethics approval

All experimental procedures with rabbits were conducted according to Chinese animal use guidelines and were approved by the Institutional Animal Care and Use Committee (IACUC).

